# Identifying Changes to RNA Editome in Major Depressive Disorder

**DOI:** 10.1101/419309

**Authors:** Saam Hasan, Shahriyar Mahdi Robbani, Tamanna Afroze, Giasuddin Ahsan, Muhammad Maqsud Hossain

**Author notes:** Corresponding author: Saam Hasan, Mailing address: Plot 15, Block B, Bashundhara, Dhaka 1229, Bangladesh.

## Abstract

Major Depressive Disorder (MDD) is one of the most significant psychiatric disorders in the world today. Its incidence is widespread in society and its heavy adverse impact on the quality of life is well documented. Previously genetic studies on MDD had identified a hereditary component of the disease as well as crediting RNA editing with a role in its development. The later due to an overexpression of a heavily edited isoform of the Serotonin 2c receptor. Here we used publicly available RNA sequence data from suicide patients diagnosed with MDD as well as controls for identifying RNA editing sites unique to MDD. After variant calling and several steps of filtering, we identified 142 unique RNA editing sites in the MDD patients. These included intronic, downstream, UTR3 and exonic edits. The latter comprising several amino acid changes in the encoded protein. The genes implicated to be uniquely edited in MDD included the aforementioned and previously implicated Serotonin 2c receptor, others involved in functions that play roles in depression and suicide such as Cannabinoid Receptor 1, Frizzled Class 3 Receptor, Neuroligin 3 and others.

## Introduction

Major Depressive Disorder represents one of the most statistically and clinically significant psychiatric disorders in the world. Its adverse impact on the quality of life has already been well documented by research (Kapp, 2001). MDD significantly reduces work capacity through its affect on the willingness to perform and work and through the physical damage it does through body function dysregulations. Its impact also extends to personal life as individuals suffering from MDD tend to struggle at maintaining and keeping up with personal relations. The disorder displays both chronic, where it persists for up to a duration three months or even more, and episodic versions which comprises of separate episodes beginning and ending independently of others. The major observable symptom of the condition is unhealthy depressed state of mind that hampers most activities as well a loss of positive emotions and pleasure towards most things in life (Patten et al., 2009). Diagnosis and identification of individuals suffering from MDD relies on the observation of symptoms in a given time span. This is taken as observing five or more symptoms associated with MDD within a span of five days. The symptoms included as a measure of MDD are as per the *Diagnostic and Statistical Manual of Mental Disorders* (Diagnostic and statistical manual of mental disorders, 2002). Research has in the past identified possible genetic factors predisposing one to the risk of developing MDD. This has identified heritable and familial factors as contributing to the development of the condition. The latter showing that genetic factors acquired through inheritance is indeed a contributing factor to the disease. This is further backed up by twin studies (Lohoff, 2010). Another kind of genetic change that has been speculated as a contributing factor albeit to a lesser degree is RNA editing. RNA editing is a form of post transcriptional change to RNA (Nishikura, 2010). It has been shown to play a role in regulation of several disease as well as increasing complexity of the transcriptome (Park et al., 2017). The only previous connection between MDD and RNA editing was the upregulation of a heavily edited isoform of the HTC2R gene which is a serotonin receptor (Lyddon et al., 2012). Although that remains a clear connection to psychiatric disorders and MDD, no specific variants or editing sites were identified specific to the condition. Here we make the first attempt at characterizing overall changes in the human RNA editome in depression and look to identify editing sites specific to MDD. We used data sequenced by Labonté et al in their study published in 2017 on sex specific transcriptional regulation in Depression (Labonté et al., 2017). We used the RNA sequence data from both MDD and Controls to compare for Editing sites found only in the MDD models. Our major goals of this study were to identify changes in editing patterns in MDD compared to control individuals. To compare the frequency of editing in the two models. And to identify specific editing sites that can serve as potential diagnostic markers for MDD and potentially suicidal behaviour.

## Methods

The data used was downloaded from NCBI SRA database using the NCBI SRA Toolkit. The prefetch and fastq-dump functions were used for this purpose (Leinonen, Sugawara and Shumway, 2010). We chose RNA sequence data from the Nucleus Acumbens region owing to its functional significance in mood and behaviour regulation (Pavuluri, Volpe and Yuen, 2017). Table 1 shows the accession numbers of the runs we used. In total five samples chosen were individuals diagnosed with MDD and committed suicide and two were controls who died naturally.

**Table 1:** Detals and SRA Accession Numbers of Data Used

| Run Number | Phenotype | Cause of Death | SRA Accession Number |
| --- | --- | --- | --- |
| 1 | MDD | Suicide | SRR5962012 |
| 3 | MDD | Suicide | SRR5961992 |
| 4 | Control | Natural | SRR5961986 |
| 5 | MDD | Suicide | SRR5961999 |
| 6 | Control | Natural | SRR5961987 |
| 12 | MDD | Suicide | SRR5961995 |
| 13 | MDD | Suicide | SRR5961996 |

We followed a modified version of the GATK best practices pipeline for variant discovery from RNA sequence data. STAR was chosen as the mapping tool. We used two pass STAR which is designed for better mapping splice junctions. We set the mismatch parameter to 3 and overhang to 49 for matching the fastq files which had 50 base pair read lengths (Dobin et al., 2012). Afterwards we used the GATK pipeline for calling variants (DePristo et al., 2011). This included started with adding read groups and marking duplicates with picard. Then we split N Cigar reads using GATK’s SplitNCigarReads functionality, recalibrated bases with BaseRecalibrator and finally called variants with HaplotypeCaller. We then split the SNPs and indels, keeping only the former using SelectVariants. Indels do not represent editing sites and hence we didn’t consider those variants any further. The variants were then analysed with the Mann-Whitney-Wilcoxon Rank Sum Test using the BaseQRankSum function of GATK’s VariantAnnotator. These were subsequently marked for common variants having dbsnp entries using VariantAnnotator again before being filtered for SNP clusters of 3 SNPs within 35 base pair windows using VariantFiltration. The variants were then annotated using Annovar developed by Wang labs. Annovar’s built in RefGene database was used for annotating variants (Wang, Li and Hakonarson, 2010). Afterwards we filtered variants for actual editing sites from SNPs in a series of steps. Through STAR’s mapping quality filters we had already filtered out most possible wrong base calls during sequencing. Now we first found the common variants between all the data from individuals with MDD who committed suicide. This followed the rationale from Ramaswami et al who established that any rare SNP is unlikely to be common between two separate individuals being studied and any common variants thus are likely to be editing sites (Ramaswami et al., 2012). Afterwards we removed any editing sites that were also found in either of the controls. Next we used the rationale followed by Zhang and Xiao that allelic linkage should not be observed in editing sites in contrast to genomic SNPs (Zhang and Xiao, 2015). We took the predicted genotype for each variant from HaplotyCaller and compared it to the observed number of alleles of reads harboring that particular variant. Variants where the genotype and allele ratio were in agreement were then discarded as possible genomic SNPs. The rest were retained and subsequently filtered again through those that did not pass the minimum z score value from the BaseQRankSum test, those having a Fisher Strand bias test score of over 30 and finally those variants with a allele depth value of under 2. Finally, variants with dbsnp entries were removed to leave the final list of RNA editing sites specific to depression. Further functional annotation and pathway analysis of these final RNA editing sites was carried using ReactomPA on R available on Bioconductor and the Database for Annotation, Visualization and Integrated Discovery (DAVID) (Huang, Sherman and Lempicki, 2009, Yu and He, 2016).

## Results

In each of the five samples from suicide victims previously diagnosed with MDD, we initially found approximately 606000, 401000, 539000, 545000, 703000 variants respectively for the runs labelled 1. 3, 5, 12, 13, including variants with dbsnp entries. The controls, labelled Run 4 and 6, contained approximately 270000 and 530000 variants respectively, including dbsnp variants. After removing common variants with dbsnp entries, the numbers decreased significantly for all the runs. Table 2 shows the exact number of variants for both with and without dbsnp entries.

**Table 2:** Number of Variants Identified in Each Run With or Without Known Variants (dbsnp)

| Run | Run1 | Run3 | Run4 | Run5 | Run6 | Run12 | Run13 |
| --- | --- | --- | --- | --- | --- | --- | --- |
| Total | 606938 | 401223 | 270214 | 599935 | 530667 | 545738 | 703541 |
| Total Without known variants (dbsnp) | 374468 | 214035 | 137447 | 329086 | 293650 | 295502 | 382241 |

Out of these we found the common positions where variants were discovered in all the MDD models. The same was done for the two control runs. We then subtracted the variants that were common to the controls from the variants common to the MDD runs as well removing known variants with dbsnp IDs. This left us with a total of 716 variants. At this stage we applied the concept for distinguishing RNA editing sites and genomic SNPs utilized by Zhang and Xiao as explained in methods. A total of 250 variants were filtered out leaving 466 RNA editing sites. These were then filtered for BaseQRankSum score, Fisher strand bias score and allele depth (methods). This narrowed down potential editing sites to 349. Finally we removed any site that occurred in either of the controls individually. The final list of RNA editing sites contained 143 editing sites. Table 3 lists these sites as well as their relevant functional information.

**Table 3:**
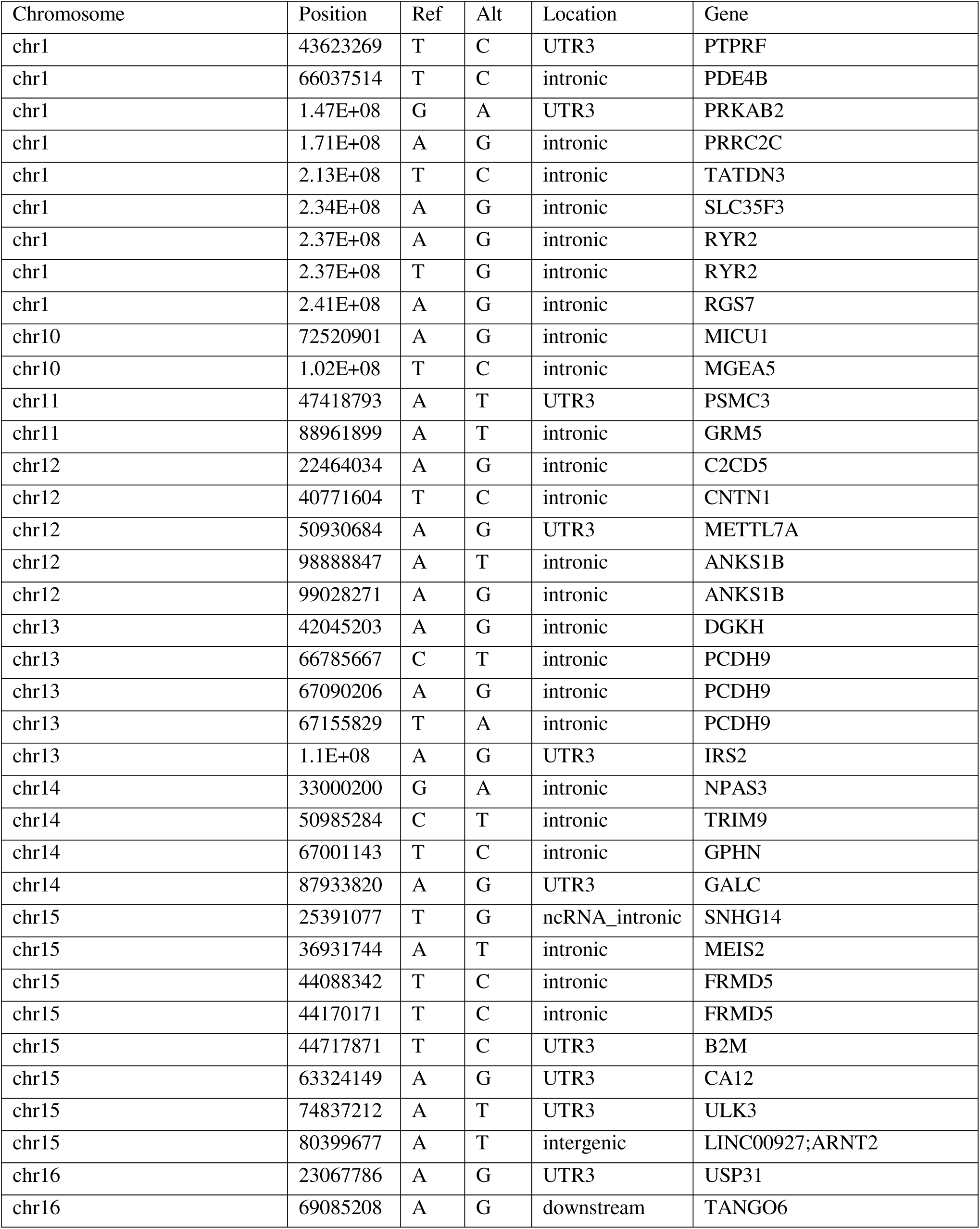
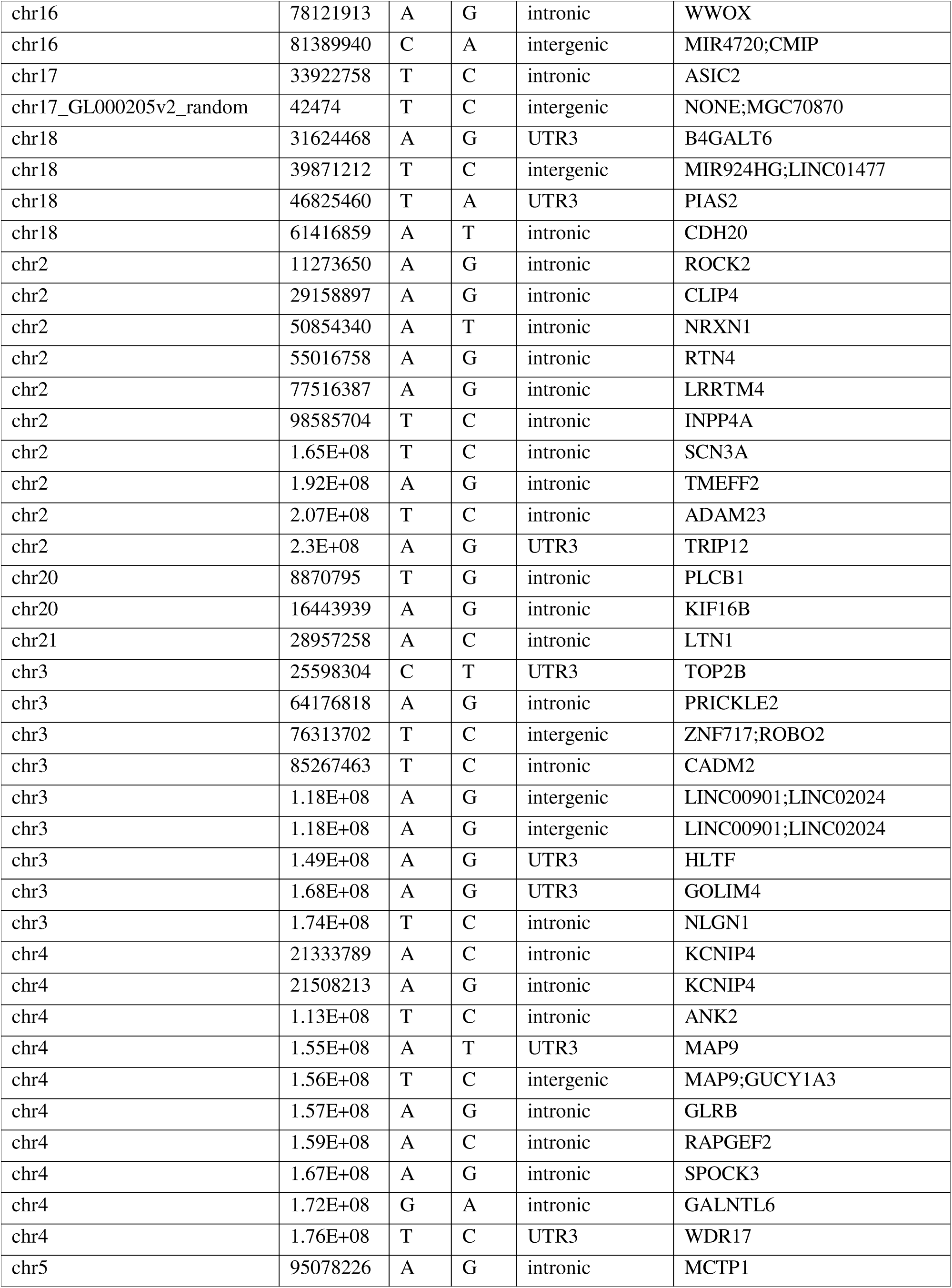
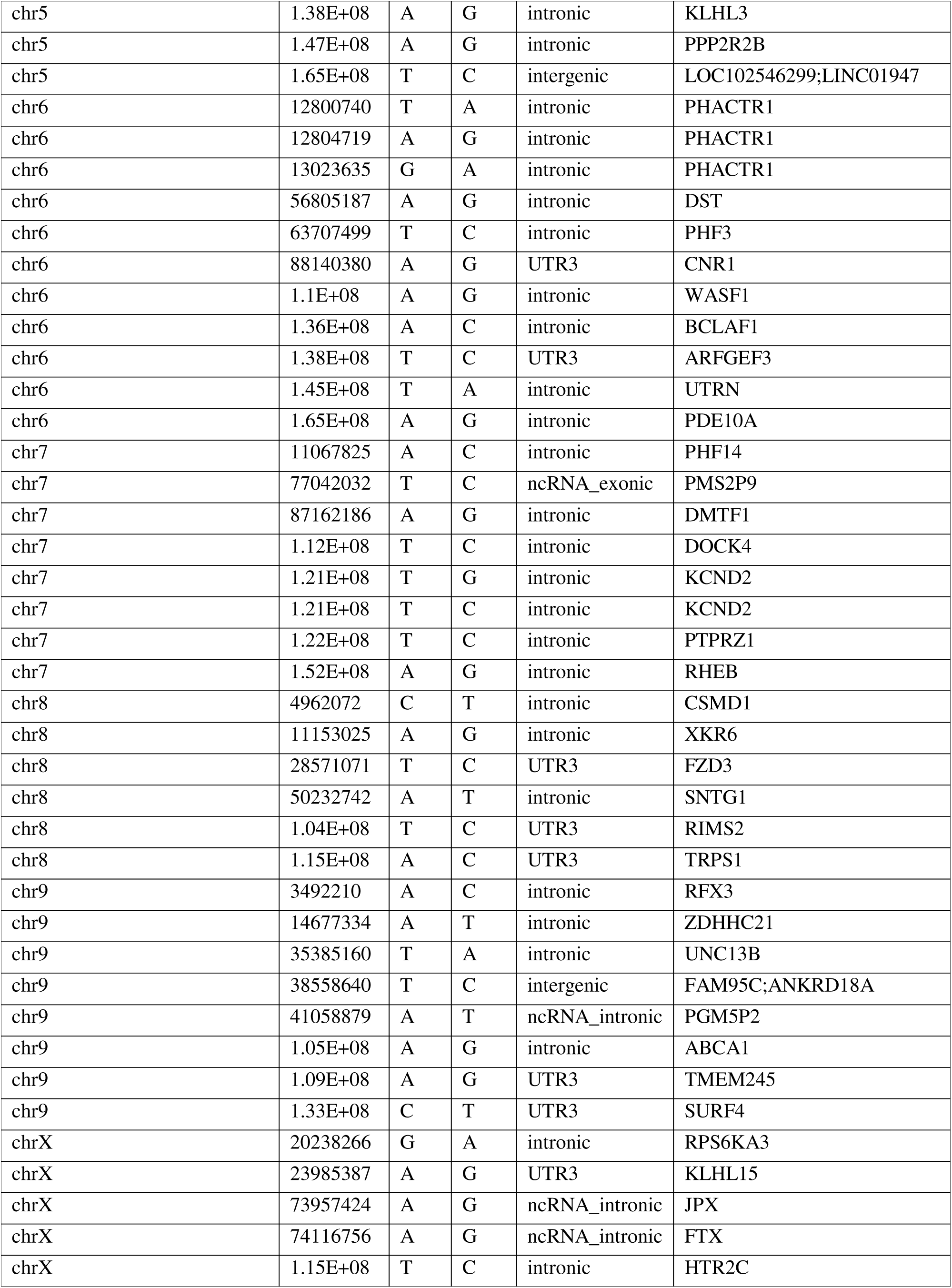
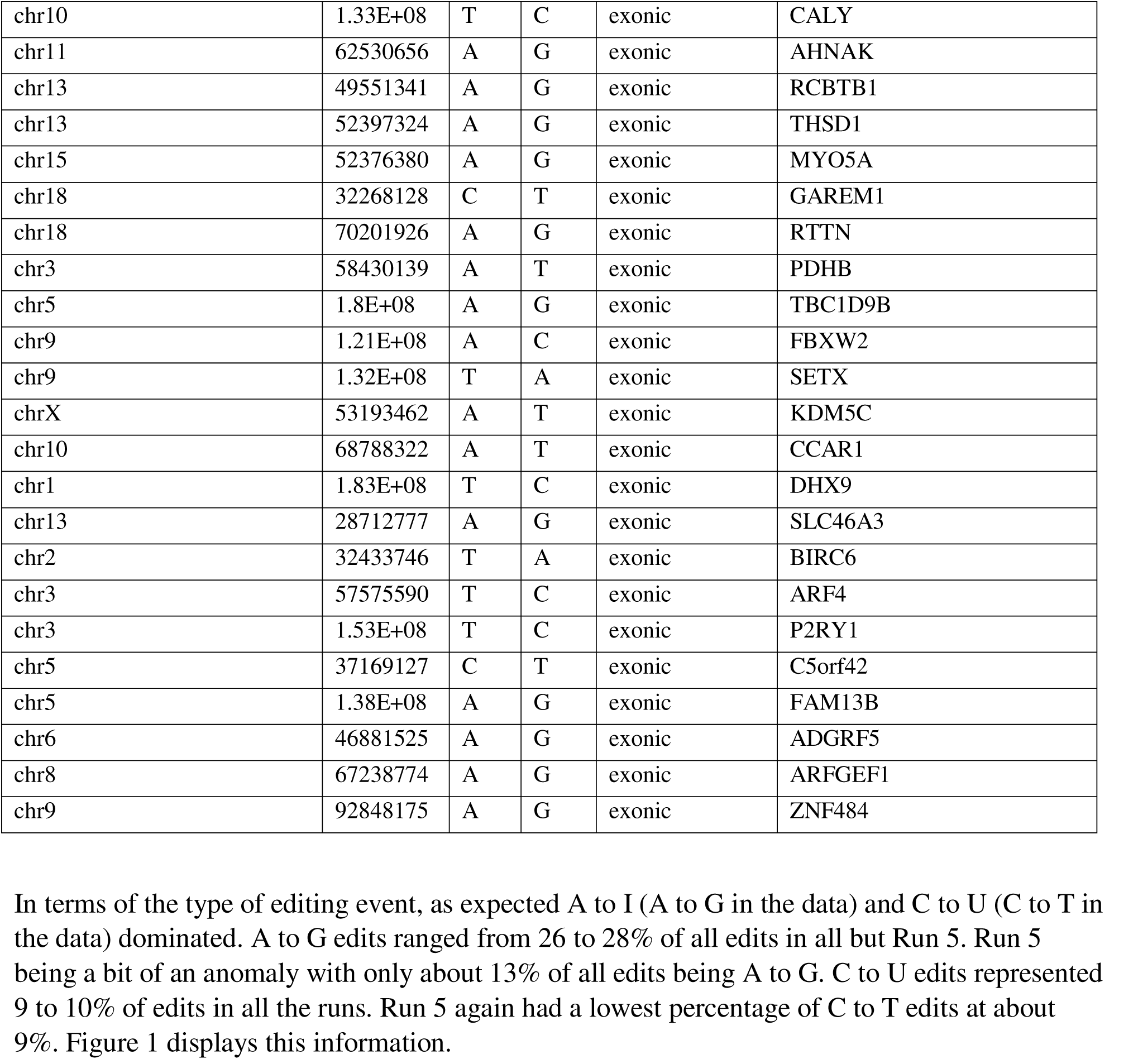
Final list of Annotated RNA Editing Sites

In terms of the type of editing event, as expected A to I (A to G in the data) and C to U (C to T in the data) dominated. A to G edits ranged from 26 to 28% of all edits in all but Run 5. Run 5 being a bit of an anomaly with only about 13% of all edits being A to G. C to U edits represented 9 to 10% of edits in all the runs. Run 5 again had a lowest percentage of C to T edits at about 9%. Figure 1 displays this information.

**Figure 1:**
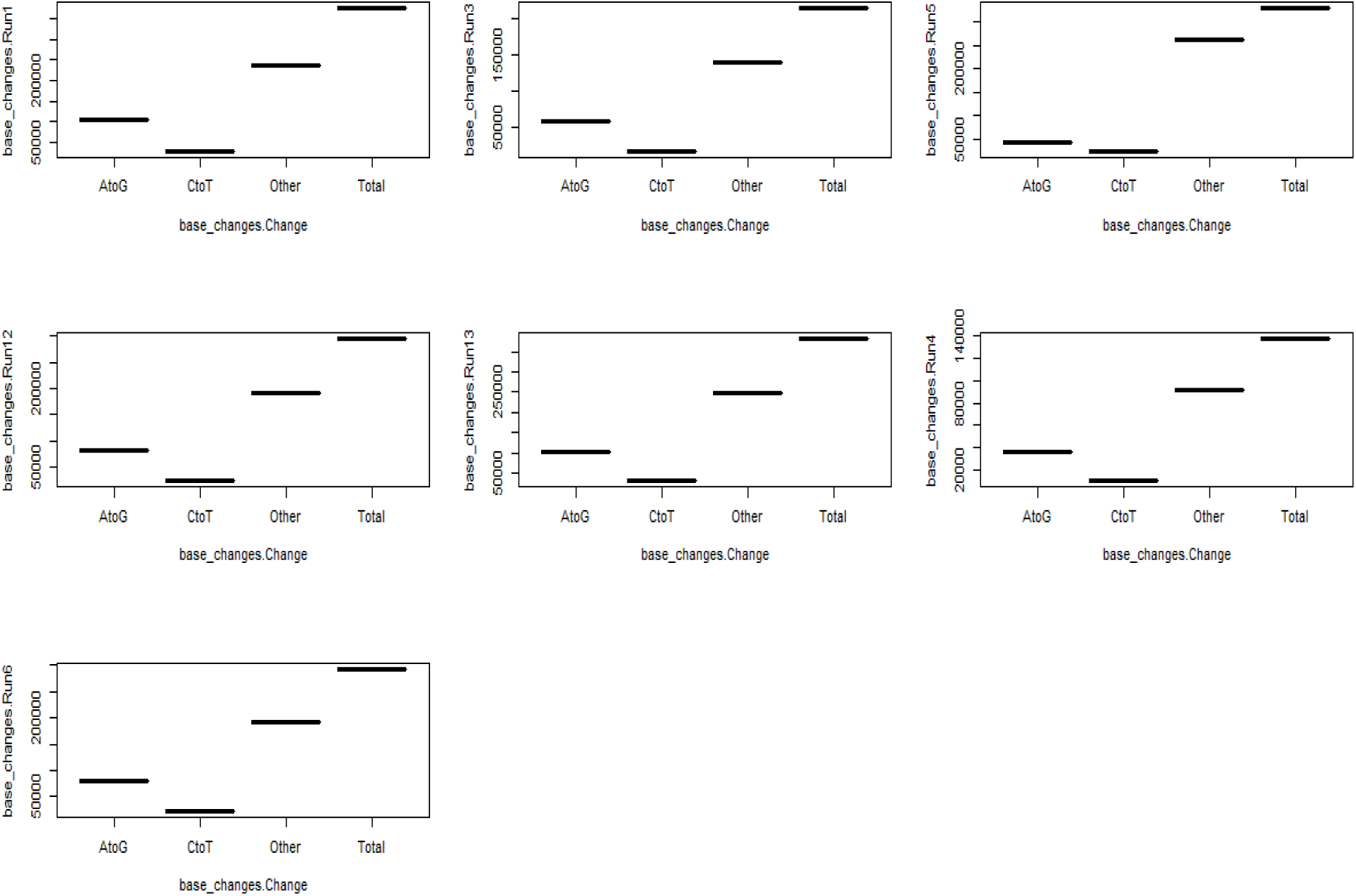
Types of Base Changes for Each Run (After Removing Known Variants)

In terms of exonic changes, which are usually more difficult to identify for RNA editing sites, we found 23 editing sites within exons in our final list of MDD specific editing sites. Among these 12 induced nonsynonymous amino acid changes in the encoded protein. Out of the amino acid changes, four were Methionine to Threonine changes. We performed a pathway enrichment analysis of the list of genes using the ReacomPA package available on Bioconductor on R. This subsequently gave us several relevant functional pathways that maybe dysregulated in MDD as a result of these editing events. Figure 2 shows a dotplot for the pathway enrichment analysis.

**Figure 2:**
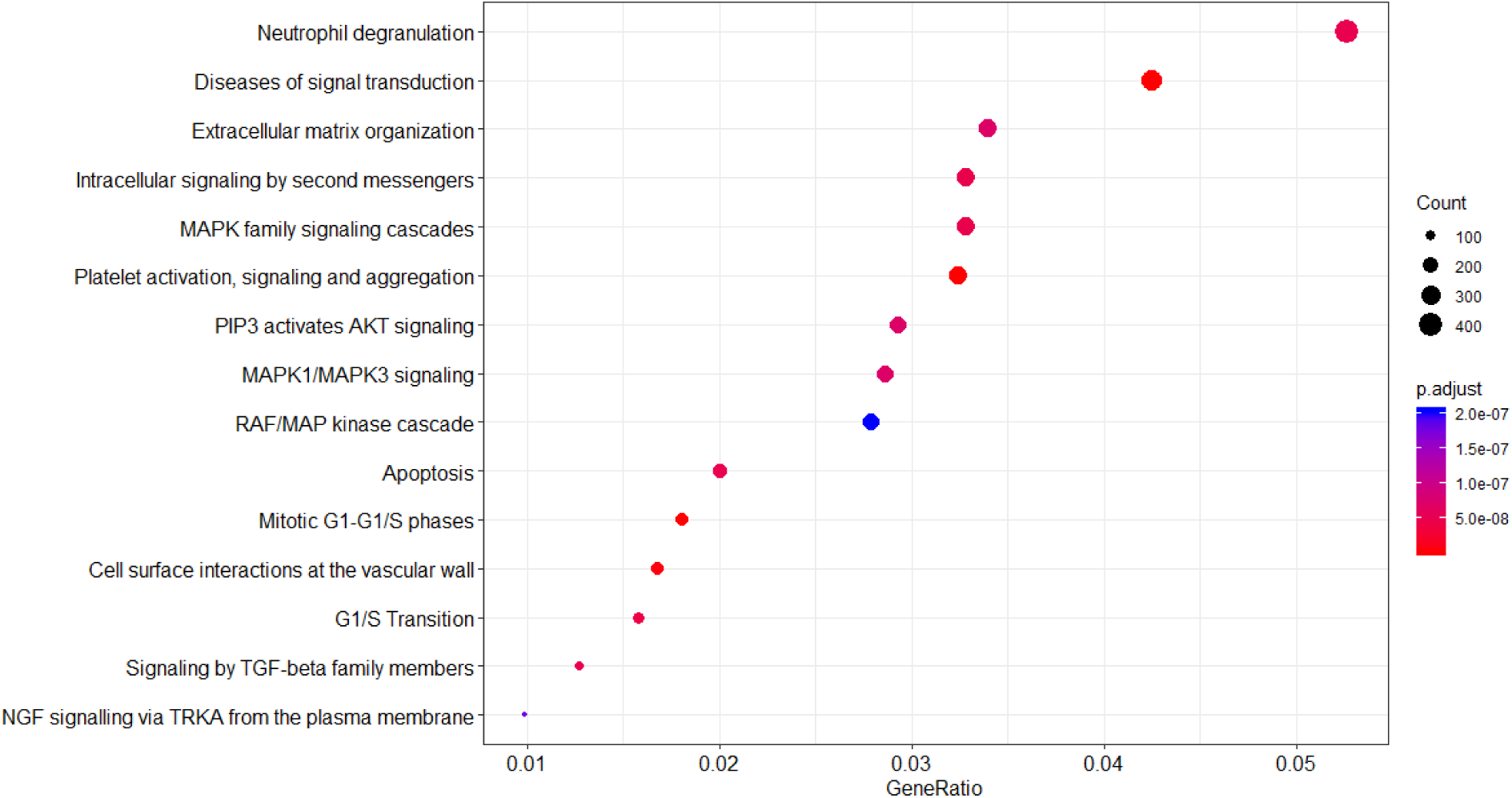
Results of Pathway Enrichment Analysis of Final List of Editing Sites. This shows the 15 most significantly affected functional pathways by the Genes Harbouring Said Editing Sites

## Discussion

We attempted to identify new RNA editing sites in individuals diagnosed with Major Depressive Disorder and who subsequently were suicide victims. We also wished to characterize any changes in RNA editome that may be specific to the disease. While we did not find any alterations to overall RNA editing pattern, we identified a number of new editing sites that were unique to the disease models. A significant number of these were exonic and even caused nonsynonymous codon changes leading to amino acid substitutions. Among these, our findings included the Serotonin 2c Receptor. However while our initial results included a large number of editing sites in this gene from the MDD models, after the filtering steps we were only left with one intronic editing site in the gene in our final list. This could possibly imply a lesser role for the edited isoform of the gene’s transcript in MDD than previously speculated. Several of our other genes however offer potential new markers for the disease. The Cannabinoid Receptor 1 has functions implicated in depression associated disorders and may offer a potential marker through the editing site we found here (an A to I edit in its UTR3). The frizzled class 3 receptor is another gene previously implicated in depression. This gene harbored a T to C edit in its UTR3. Neuroligin 3 was another gene we found to have MDD specific editing that is involved in depression associated pathways. This contained an intronic T to C editing site. Lastly phosphodiesterase 4B was our fifth gene to have a direction functional relationship with depression. This contained an A to I intronic edit. In addition to these, several of the genes we identified have functions associated with bi polar disorder such as multiple C2 and transmembrane domain containing 1, Phosphodiesterase 10A and aryl hydrocarbon receptor nuclear translocator 2. We also found several genes implicated in schizophrenia that contained unique RNA editing sites. This included protein phosphatase 2 regulatory subunit B’beta, protein tyrosine phosphatase, receptor type Z1, reticulon 4 and others. Other genes we identified were relevant in context of mental retardation, attention deficiency, autism, smoking disorder and alcoholism (UniProt Consortium, 2018). All of which in terms of their negative impact on quality of life can have potential contributions to depression and suicidal behavior. At the time of submission of this paper, we are still furthering this research through running analysis on more RNA sequence samples from MDD diagnosed suicide patients, controls and non MDD suicide patients. Here we present our initial findings that may serve as potential markers for the identification and early diagnosis of MDD and potential suicidal tendencies.

## References

DePristo, M., Banks, E., Poplin, R., Garimella, K., Maguire, J., Hartl, C., Philippakis, A., del Angel, G., Rivas, M., Hanna, M., McKenna, A., Fennell, T., Kernytsky, A., Sivachenko, A., Cibulskis, K., Gabriel, S., Altshuler, D. and Daly, M. (2011). A framework for variation discovery and genotyping using next-generation DNA sequencing data. Nature Genetics, 43(5), pp.491–498.

Diagnostic and statistical manual of mental disorders. (2002). Washington, DC: American Psychiatric Association.

Dobin, A., Davis, C., Schlesinger, F., Drenkow, J., Zaleski, C., Jha, S., Batut, P., Chaisson, M. and Gingeras, T. (2012). STAR: ultrafast universal RNA-seq aligner. Bioinformatics, 29(1), pp.15–21.

Huang, D., Sherman, B. and Lempicki, R. (2009). Systematic and integrative analysis of large gene lists using DAVID bioinformatics resources. Nature Protocols, 4(1), pp.44–57.

Kapp, C. (2001). WHO report aims to increase understanding of mental health. The Lancet, 358(9289), p.1248.

Labonté, B., Engmann, O., Purushothaman, I., Menard, C., Wang, J., Tan, C., Scarpa, J., Moy, G., Loh, Y., Cahill, M., Lorsch, Z., Hamilton, P., Calipari, E., Hodes, G., Issler, O., Kronman, H., Pfau, M., Obradovic, A., Dong, Y., Neve, R., Russo, S., Kazarskis, A., Tamminga, C., Mechawar, N., Turecki, G., Zhang, B., Shen, L. and Nestler, E. (2017). Sex-specific transcriptional signatures in human depression. Nature Medicine, 23(9), pp.1102–1111.

Leinonen, R., Sugawara, H. and Shumway, M. (2010). The Sequence Read Archive. Nucleic Acids Research, 39(Database), pp.D19–D21.

Lohoff, F. (2010). Overview of the Genetics of Major Depressive Disorder. Current Psychiatry Reports, 12(6), pp.539–546.

Lyddon, R., Dwork, A., Keddache, M., Siever, L. and Dracheva, S. (2012). Serotonin 2c receptor RNA editing in major depression and suicide. The World Journal of Biological Psychiatry, 14(8), pp.590–601.

Nishikura, K. (2012). Functions and Regulation of RNA Editing by ADAR Deaminases. Annual Review of Biochemistry, 79(1), pp.321–349.

Park, E., Guo, J., Shen, S., Demirdjian, L., Wu, Y., Lin, L. and Xing, Y. (2017). Population and allelic variation of A-to-I RNA editing in human transcriptomes. Genome Biology, 18(1).

Patten, S., Kennedy, S., Lam, R., O’Donovan, C., Filteau, M., Parikh, S. and Ravindran, A. (2009). Canadian Network for Mood and Anxiety Treatments (CANMAT) Clinical Guidelines for the Management of Major Depressive Disorder in Adults. I. Classification, Burden and Principles of Management. Journal of Affective Disorders, 117, pp.S5–S14.

Pavuluri, M., Volpe, K. and Yuen, A. (2017). Nucleus Accumbens and Its Role in Reward and Emotional Circuitry: A Potential Hot Mess in Substance Use and Emotional Disorders. AIMS Neuroscience, 4(1), pp.52–70.

Ramaswami, G., Lin, W., Piskol, R., Tan, M., Davis, C. and Li, J. (2012). Accurate identification of human Alu and non-Alu RNA editing sites. Nature Methods, 9(6), pp.579–581.

UniProt Consortium, T. (2018). UniProt: the universal protein knowledgebase. Nucleic Acids Research, 46(5), pp.2699–2699.

Wang, K., Li, M. and Hakonarson, H. (2010). ANNOVAR: functional annotation of genetic variants from high-throughput sequencing data. Nucleic Acids Research, 38(16), pp.e164–e164.

Yu, G. and He, Q. (2016). ReactomePA: an R/Bioconductor package for reactome pathway analysis and visualization. Molecular BioSystems, 12(2), pp.477–479.

Zhang, Q. and Xiao, X. (2015). Genome sequence–independent identification of RNA editing sites. Nature Methods, 12(4), pp.347–350.

